## Supplementary material for "The effect of gravity on hand spatio-temporal kinematic features during functional movements": Supplemetary analyses on gravity-related kinematic performances of healthy behaviour during a functional task.

### Movements Within Gravity Condition

Examining each movement (M1-M9), both absolute hand positions and velocities exhibit qualitative similarities across various subjects and trials.

The initial hypothesis suggests that when performing movements in the same direction with respect to gravitational force, the proposed kinematic features remain unaffected by the starting and target positions of the three objects.

#### *Movement Time*

As for MT, Friedman test did not present statistical significance for $g^-$ did not show a significant effect (p = 0.32, $\chi^2(2) = 2.28$), meanwhile for $g^+$ it did show statistical significance as p = 0.003, $\chi^2(2) = 11.6$. However, Mann-Whitney tests performed over M4-M5, M4-M6, and M5-M6 resulted in non-statistically significant differences, showing p = 0.12, p = 0.13, and p = 0.94 respectively. Such result might be due to the lack of statistical power as groups considered are characterized by small number of samples (72 for each group). For $g^0$ condition, Friedman test calculated over M7, M8, and M9 presented a statistically significant effect (p = 0.0005, $\chi^2(2) = 15$). Pairwise tests performed over M7-M8 and M7-M9 resulted in a statistically significant difference (p = 0.006 and p = 0.0096 respectively) meanwhile M8-M9 did not result in any statistical significance (p = 0.83).

#### *Percent Time to Peak Velocity*

PTPV metric across each subject and repetition presented overall a normal distribution, therefore parametric tests were employed. For $g^-$ condition, repeated measures ANOVA test calculated over M1, M2, and M3 did not show any statistically significant effect (p = 0.06, $F(2)(2) = 3.9$). Same effect was observed for $g^+$ (repeated measures ANOVA test calculated over M4, M5, and M6 did not present a statistically significant effect (p = 0.06, $F(2)(2) = 2.93$)). For $g^0$ condition, repeated measures ANOVA test calculated over M7, M8, and M9 presented a statistically significant effect ($p < 0.0001$, $F(2) = 15$). Pairwise tests performed over M7-M9 resulted in a statistical significance ($p < 0.0001$), M7-M8 presented statistical significance (p = 0.018), and M8-M9 a non statistically significant difference (p = 0.03).

#### *Percent Time to Peak Standard Deviation*

PTPSD metric across each subject and repetition presented overall a normal distribution, therefore parametric tests were employed.

For $g^-$ condition, repeated measures ANOVA test calculated over M1, M2, and M3 did not show any statistically significant effect (p = 0.58, $F(2) = 0.55$). Same effect was observed for $g^+$ condition (repeated measures ANOVA test calculated over M4, M5, and M6 did not present a statistically significant difference (p = 0.76, $F(2) = 0.27$)) and $g^0$ (repeated measures ANOVA test calculated over M7, M8, and M9 did not present a statistically significant effect (p = 0.87, $F(2) = 0.14$)).

#### *Spatial Arc Length*

SPARC metric across each subject and repetition presented overall a normal distribution, therefore parametric tests were employed. Repeated measures ANOVA tests did not yield any statistical significance for $g^-$ (p = 0.95, $F(2) = 0.05$), $g^+$ (p = 0.39, $F(2) = 0.94$), and $g^0$ (p = 0.33, $F(2) = 2.43$).

#### *General Conclusion*

96% (104 out of 108) of pair-wise post-hoc comparisons performed among MT, PTPV, PTPSD, and SPARC metrics did not result in any statistical significance (p<0.017) therefore the previously posited hypothesis is not rejected.

### Between subjects analysis

Following results obtained from the previous section, movements M1-M3, M4-M6, and M7-M9 were pooled together as representative of one single movement for $g^-$, $g^+$, and $g^0$ repeated 9 times. Based on this theoretical assumption the variability of metrics NVP, MT, PTPSD, PTPV, and SPARC was studied between subjects. Figures 1 and 3 report box-charts of each metric for the 24 subjects recruited (in blue) and the total distribution (in red) for $g^-$, $g^+$, and $g^0$. Statistical tests were conducted to assess the effect of recruited subjects on the interested index and, if present, pair-wise post-hoc tests were employed to compare each subject's distribution to the overall one.

#### *Movement Time and Number of Velocity Peaks*

Friedman test was performed to investigate the effect of different subjects on MT. As statistical significance was found for all three conditions $g^-$, $g^+$, and $g^0$ (p<0.0001), Mann-Whitney post-hoc tests were conducted. Subjects 5, 6 and 16 constantly showed $p < 0.0001$ as they presented different duration of the movement with respect to the global median. Test on subject 10 resulted in $p < 0.001$ and $p < 0.0001$ for $g^-$ and $g^0$ respectively. Subject 11 and 22 only presented $p < 0.001$ for $g^+$, subject 13 $p < 0.001$ in $g^0$, subject 18 $p < 0.001$ in $g^+$, and subject 22 $p < 0.001$ both in $g^+$ and $g^0$ (Figure 1A). Friedman test on

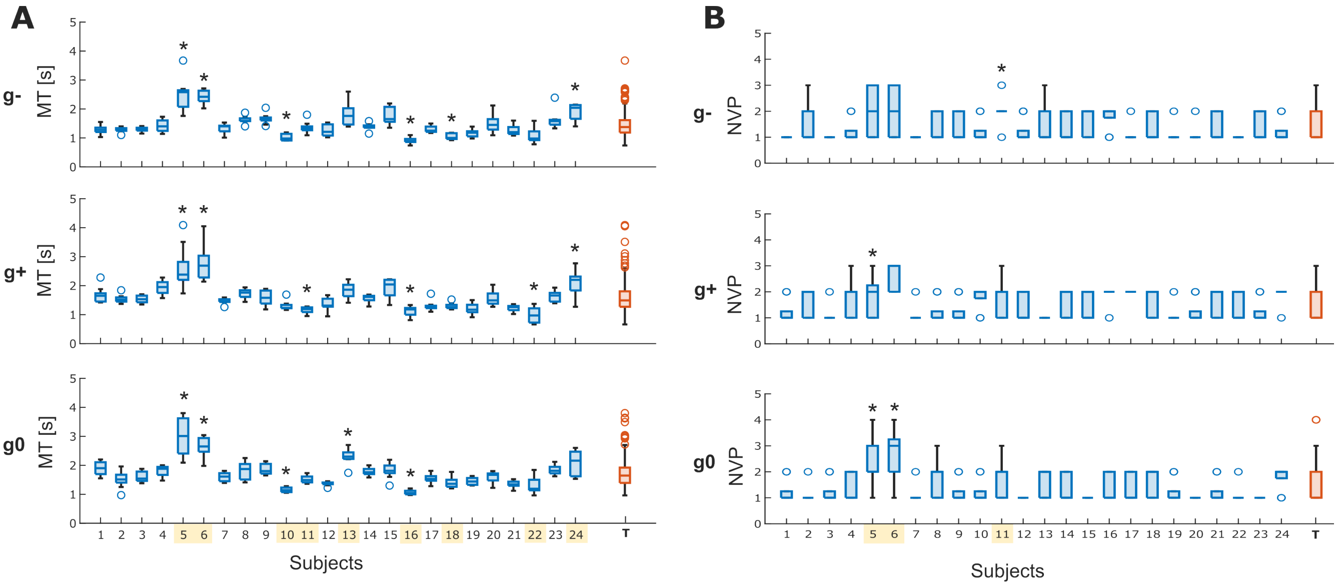


**Figure 1. Metrics calculated for each subject** Rows of the graph represents boxplots for each subject (in blue) and a boxplot for the total distribution (in red) for $g^{-,+,0}$. A) Related to MT, B) related to NVP. * indicates statistical significance ($p < 0.05$) between the interested subject and the total distribution.

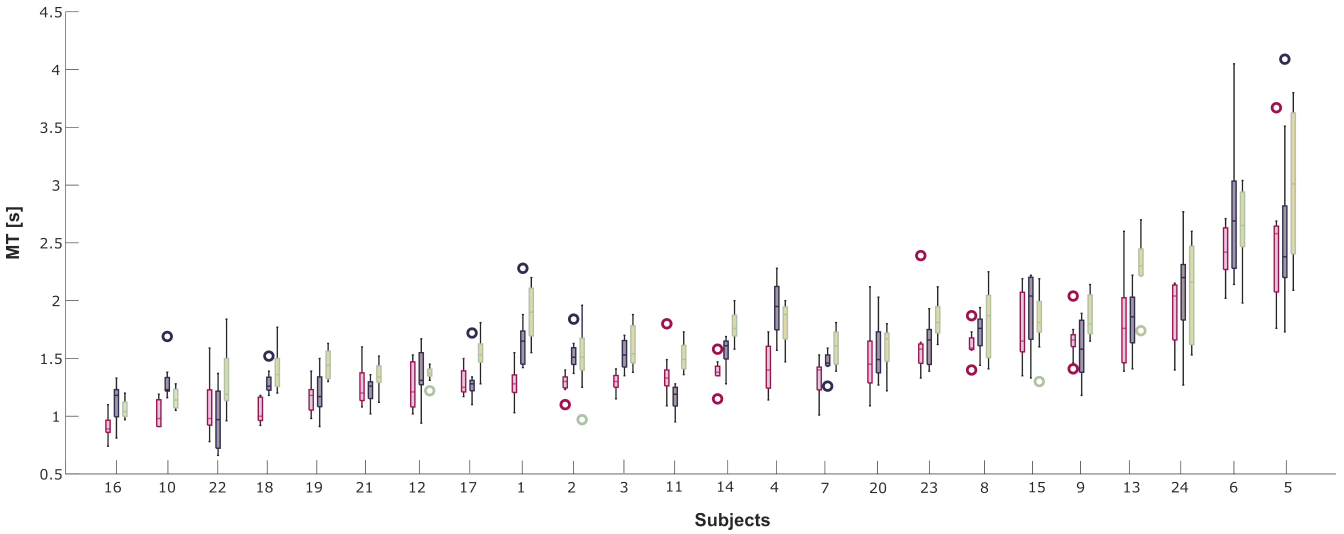


**Figure 2. Distribution of MT for direction-dependent set of movements for each subject, sorted in increasing order with respect to $g^-$** Purple is related to $g^-$ movements, blue to $g^+$, and green to $g^0$.

NVP yielded a statistical significance ($p < 0.0001$) for the three sets of movement therefore Mann-Whitney pair-wise test was conducted. Subject 5 presented $p < 0.0001$ for $g^0$, subject 6 $p < 0.0001$ for $g^{-,0}$ and subject 11 $p < 0.0001$ for $g^+$ (Figure 1B).

As 9 subjects out of 24 presented behaviour different from the total distribution for either $g^-$, $g^+$, or $g^0$, further analysis was conducted to investigate whether subjects tended to perform coherently slower or faster across different direction-dependent trajectories. Figure 2 presents subjects sorted in increasing MT for $g^-$ (in purple), with subject 16 being the fastest and subject 5 the slowest. The same order was applied to $g^+$ (in blue) and $g^0$ (in green). Even though neutral movements ($g^0$) tend to be slower than the ones against gravity ($g^-$) or propelled by gravity ($g^+$), each subject seems to apply comparable timing within sets but different between other individuals recruited. To exclude any dependency of the duration of movement from individuals' biomechanical constraints, MT was also normalized with respect to the each subject's upper-limb length (arm and forearm). As results did not change, the influence of different arm percentiles to MT is excluded.

Overall results suggest that a central measurement of MT can't be derived from the total dataset of the proposed pick-and-place task, as behaviour is different between subjects.

#### Percent Time to Peak Velocity, Percent Time to Peak Standard Deviation, and Spatial Arc Length

Repeated measures ANOVA for PTPV, PTPSD, and SPARC did not result in any statistical significance for each of the direction-dependent sets of movements. Particularly, for PTPV p-values were 0.11, 0.55, and 0.61 respectively for $g^{-}$, $g^{+}$, and $g^{0}$ (Figure 3A). Concerning PTPSD, p-values were 0.69, 0.13, and 0.81 respectively (Figure 3B). Lastly, for SPARC p-values were 0.88, 0.55, and 0.23 respectively (Figure 3C).

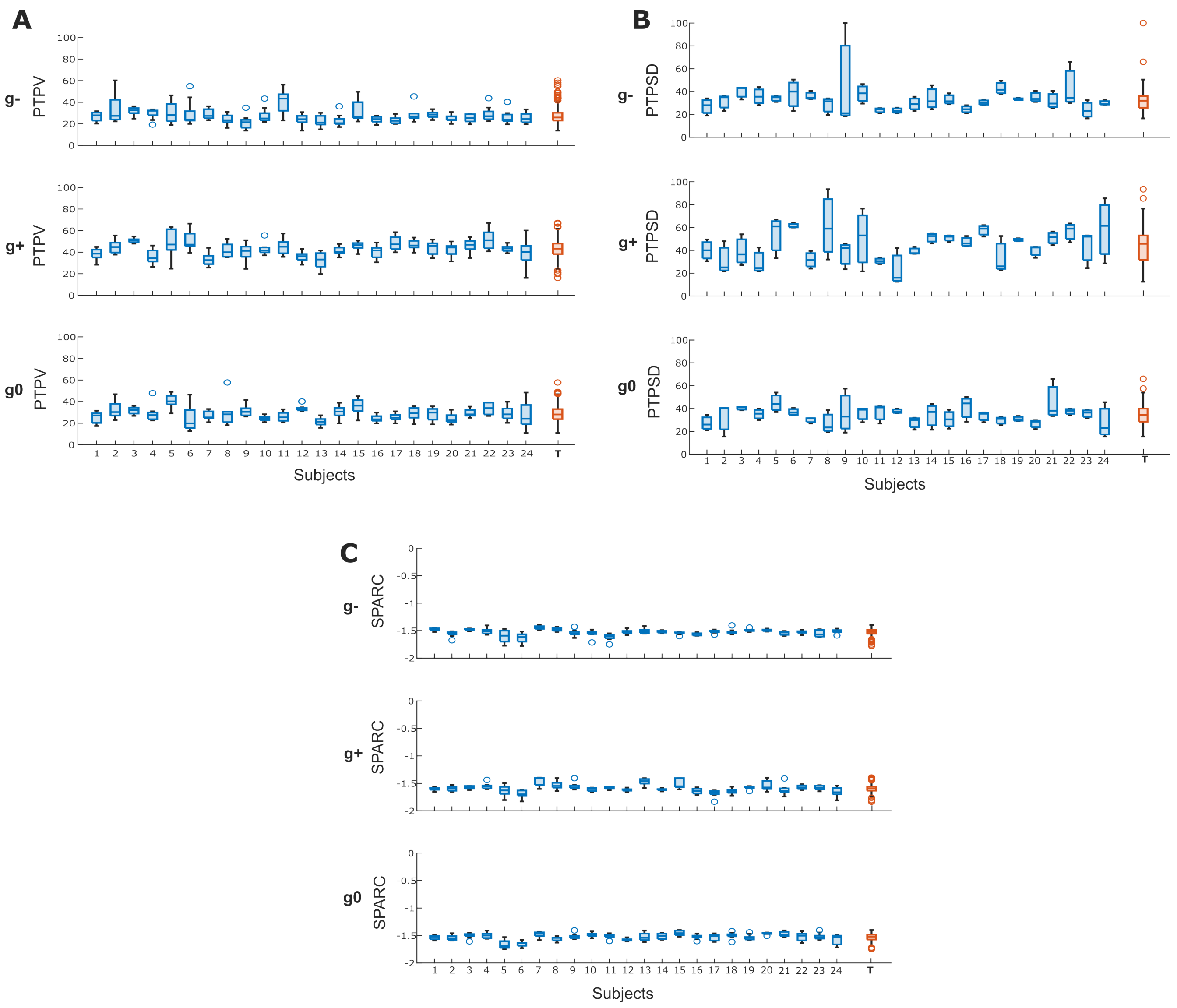


**Figure 3. Metrics calculated for each subject** Rows of the graph represents boxplots for each subject (in blue) and a boxplot for the total distribution (in red) for $g^{-,+,0}$. A) Related to PTPV, B) related to PTPSD, and C) related to SPARC).

#### General Conclusion

MT was the only metric in which few subjects (5, 6, and 16) presented consistent behaviour different from the overall distribution. As the task was self-paced, subjects did not present a common Movement Time. Such result suggests that the duration of movement (MT) is uncorrelated to the selected metrics but NVP. On the contrary, PTPV, PTPSD, and SPARC presented a common behaviour across subjects for the same movement direction.

Figure 4 shows the relationship between metrics across subjects and repetitions. PTPSD was not reported as it represents

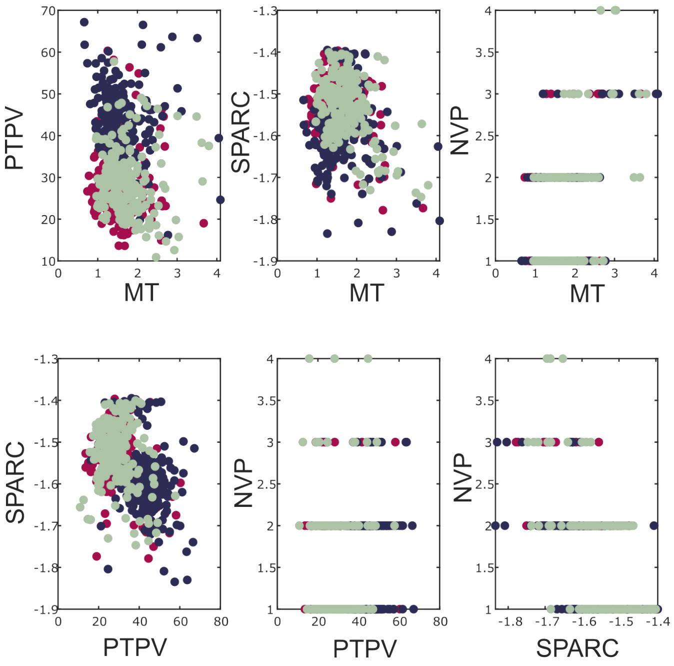


**Figure 4.** **Relationship between metrics** MT, SPARC, PTPV and NVP were correlated between each other to check whether any remarkable trend is present for $g^-$(in purple), $g^+$ (in blue), and $g^0$ (in green).

the averaged standard deviation across the 9 repetitions for each subject. As it's possible to notice, distributions related to *MT versus PTPV* and *MT versus SPARC* do not present a geometrical trend but rather a chaotic relationship. The same observation is found for *SPARC versus PTPV*. Concerning the graph MT-NVP, a higher duration of movements leads to an increase in NVP, which was expected as slower movements are associated to less smooth trajectories and multiple peaks in the velocity profiles. A similar outcome is found for NVP-SPARC relationship as less negative numbers of SPARC are associated to smoother movements.

Overall, even though MT is subject-dependent the remaining kinematic metrics do not present same non-linear behaviour across subjects, stressing how the proposed metrics but NVP are not influenced by duration of the movement.
